# Propofol anesthesia destabilizes neural dynamics across cortex

**DOI:** 10.1101/2023.11.24.568595

**Authors:** Adam J. Eisen, Leo Kozachkov, Andre M. Bastos, Jacob A. Donoghue, Meredith K. Mahnke, Scott L. Brincat, Sarthak Chandra, Emery N. Brown, Ila R. Fiete, Earl K. Miller

## Abstract

Every day, hundreds of thousands of people undergo general anesthesia. One hypothesis is that anesthesia disrupts dynamic stability, the ability of the brain to balance excitability with the need to be stable and thus controllable. We tested this hypothesis using a new method for quantifying population-level dynamic stability in complex systems, **De**layed **L**inear **A**nalysis for **S**tability **E**stimation (**DeLASE**). Propofol was used to transition animals between the awake state and anesthetized unconsciousness. DeLASE was applied to macaque cortex local field potentials (LFPs). We found that neural dynamics were more unstable in unconsciousness compared to the awake state. Cortical trajectories mirrored predictions from destabilized linear systems. We mimicked the effect of propofol in simulated neural networks by increasing inhibitory tone. Paradoxically, increased inhibition also destabilized the networks. Our results suggest that anesthesia disrupts dynamical stability that is required for consciousness.

## Introduction

The pharmacological action and neurophysiological response of propofol is well understood, but how it renders unconsciousness is not. Propofol boosts inhibition through GABA_A_ receptors and significantly alters cortical dynamics^1–6^. This could disrupt the cortical communication on which consciousness depends^1,2,7^, but the exact link to theories of consciousness is not clear. Many theories of consciousness have focused on the representation and network structure involved in integrating information or linking together cortical representations^8–14^. For example, one prominent theory of consciousness posits that awareness follows from an “ignition” that produces widespread cortical spiking much like a few claps can lead to a whole audience applauding^10,15,16^. However, overly excitable and unstable states are uncontrollable, indicative of pathological conditions^17,18^. Thus, we hypothesize that a key factor in consciousness is *dynamic stability*. Brain states should be sufficiently excitable for generation of widespread activity and information integration. But they also need to be controllable and stable, reliably producing the same computations^19–23^.

Stability has long been known to be critical for brain function but early computational work investigated it in the context of convergence to a single state, involving a kind of “freezing” of neural activity^24–26^. However, normal neural activity is rarely so stationary; rather it constantly evolves through dynamic trajectories^27,28^. Thus, stability, and hence consciousness, needs to be understood in terms of a dynamic brain^21,22^. Here, we approach the analysis of anesthetic unconsciousness through the lens of *dynamic stability* (henceforth stability) (Fig. 1a), a fundamental concept in dynamical systems theory and control. Essentially, dynamic stability is a measure of the robustness of a dynamical system. The system needs to be able to recover from disturbances (e.g., distractions, random fluctuations in activity) to its normal state.

**Figure 1.**
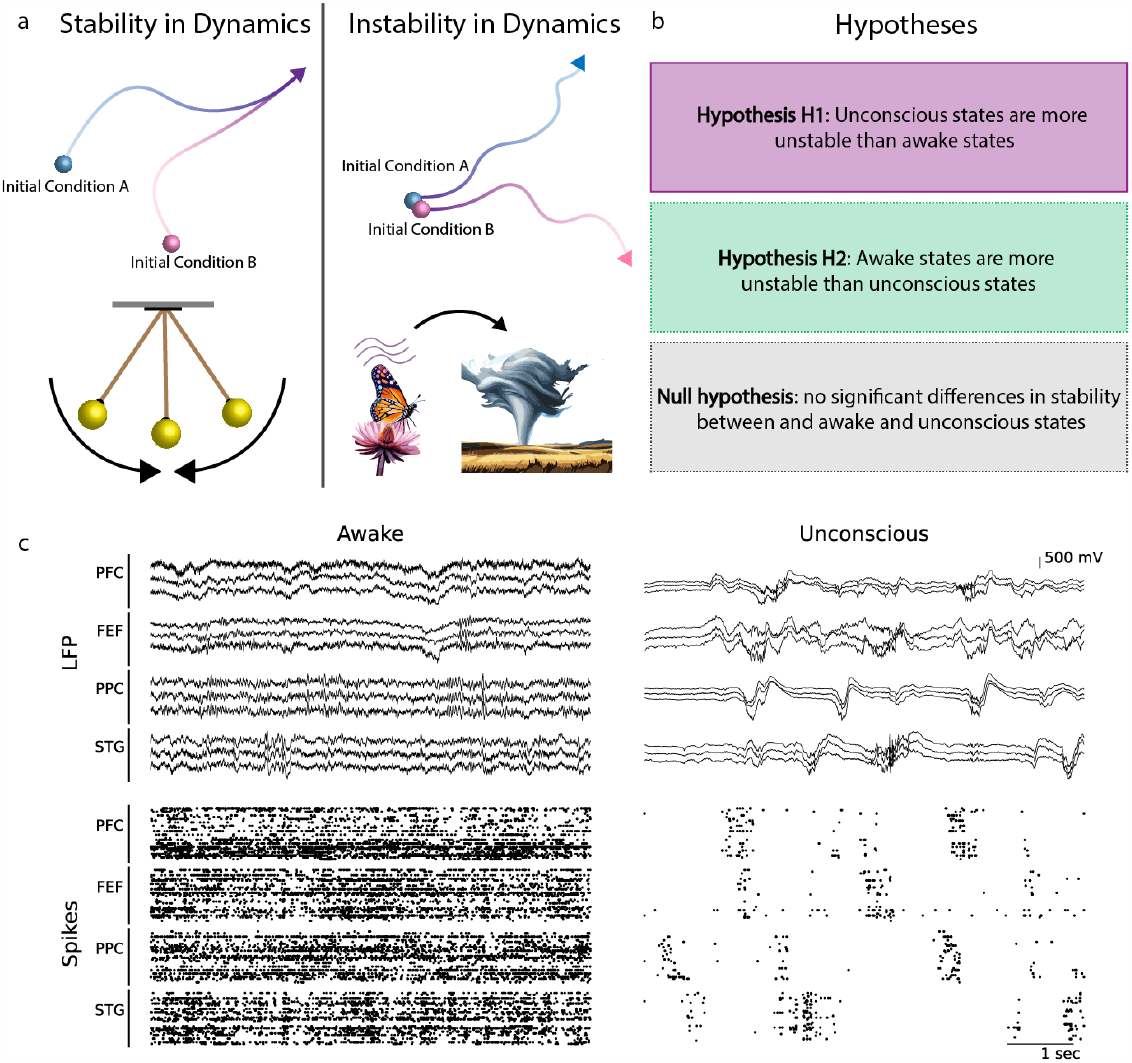
Stability and instability, and hypothesis candidates. (a) (left, top) A depiction of stability in dynamics. Starting from two distinct initial conditions, trajectories of the system converge to be the same. (left, bottom) A diagram of a stable system - a pendulum with friction. Dropped from anywhere, the pendulum will eventually converge to the bottom position. (right, top) A depiction of instability in dynamics. Starting from two very similar initial conditions, the trajectories of the system nonetheless diverge to very distinct paths. (right, bottom) A cartoon of an unstable system - the weather. A very small perturbation, the flap of a butterfly’s wings, may cause a large-scale change to the system in the future, such as a tornado^33^. (b) Three hypotheses regarding the impact of propofol anesthesia on neural dynamics. Dynamics can either be more unstable than awake dynamics, more stable than awake dynamics, or can show no significant change. (c) Sample neural data from the propofol dataset is shown. (left, top) 10 seconds of LFPs from all four brain areas during the awake. (left, bottom) Spike rasters from 10 seconds of recording from all four brain areas during the awake state. (right, top) 10 seconds of LFPs from all four brain areas during the unconscious state. (right, bottom) Spike rasters from 10 seconds of recording from all four brain areas during the unconscious state. During the awake state, LFP signals are lower amplitude and display higher frequency activity. Spiking is consistent and shows no clear coordinated bursting across areas. In the unconscious state, by contrast, LFP signals are much higher amplitude and display strong low frequency activity. This low frequency activity is strongly coordinated across areas, a phenomenon that has been hypothesized to underlie the loss of consciousness^1,2^. Spiking during the unconscious state shows clear up-state/down-state bursting patterns and is coordinated across areas.

Previous work on cortical stability during anesthesia has produced contradictory results, suggesting that anesthesia either destabilizes^29,30^ or excessively stabilizes^31,32^ neural dynamics (Fig. 1b). This could be due to a paucity of studies using high-density intracortical electrophysiology and the inability to therefore apply sufficiently rich dynamical tools to assess stability. Thus, we used a dataset of local field potential (LFP) recordings with hundreds of electrodes from multiple brain regions in two non-human primates (NHPs, specifically adult rhesus macaque monkeys) as they lost and regained consciousness due to propofol anesthesia (Fig. 1c).

We introduce a new approach - Delayed Linear Analysis for Stability Estimation (DeLASE). DeLASE directly quantifies stability in neural data. We show that this method produces high-quality models of nonlinear circuit dynamics while maintaining the simplicity and tractability of a linear dynamical system. We validated the model’s estimates of changes in dynamic stability in systems for which the ground truth stability is known. We found that propofol-induced unconsciousness is associated with destabilized neural dynamics.

## Results

### Dynamical systems approach: DeLASE method

There are three major challenges to a dynamical systems analysis of neural data.

a. **Partial observation:** neural dynamics are high dimensional, and any one sample of neural data can only hope to capture a few of the dimensions of the overall system. This leads to a very restricted partial observation.
b. **Tractability:** despite limited brain state sampling, these partial observations are high dimensional. Electrophysiology can involve hundreds of electrodes and a high LFP sampling rate, generating thousands of neural data points within seconds. Therefore, modeling neural dynamics requires a model that can reconstruct high-dimensional dynamics from partial observations and a robust, tractable, and scalable model-fitting procedure to handle vast amounts of data.
c. **Nonlinearity:** neural dynamics are complex and nonlinear, which presents difficulties for both modeling and stability analysis.

In this section, we propose a new method that leverages advances in data-driven dynamical systems analysis with strong theoretical guarantees, to tractably obtain a robust estimate of stability in partially observed nonlinear dynamical systems with a large number of variables.

### Partial observation

Takens’ Delay Embedding Theorem is a powerful result from dynamical systems theory which shows how the full attractor of a dynamical system can be reconstructed from partial observation of data by appending lagged copies of the subset of observed variables onto the existing measurements^34^. In other words, remarkably, it is possible to trade space (the set of all variables in the system) for time (multiple time observations of a subset of variables) to fully characterize a nonlinear dynamical system. As an example of this “delay embedding” principle^35–46^, the famous “butterfly attractor”^33^ of the three-dimensional Lorenz system can be reconstructed from a single observation (Fig. 2a). The key takeaway is that information is gained through considering the trajectory history of a system, in conjunction with its current state. As we hope to convey in this paper, this has tremendous implications for neural data, which is likely capturing a very small fraction of the information contained in the full neural system.

**Figure 2.**
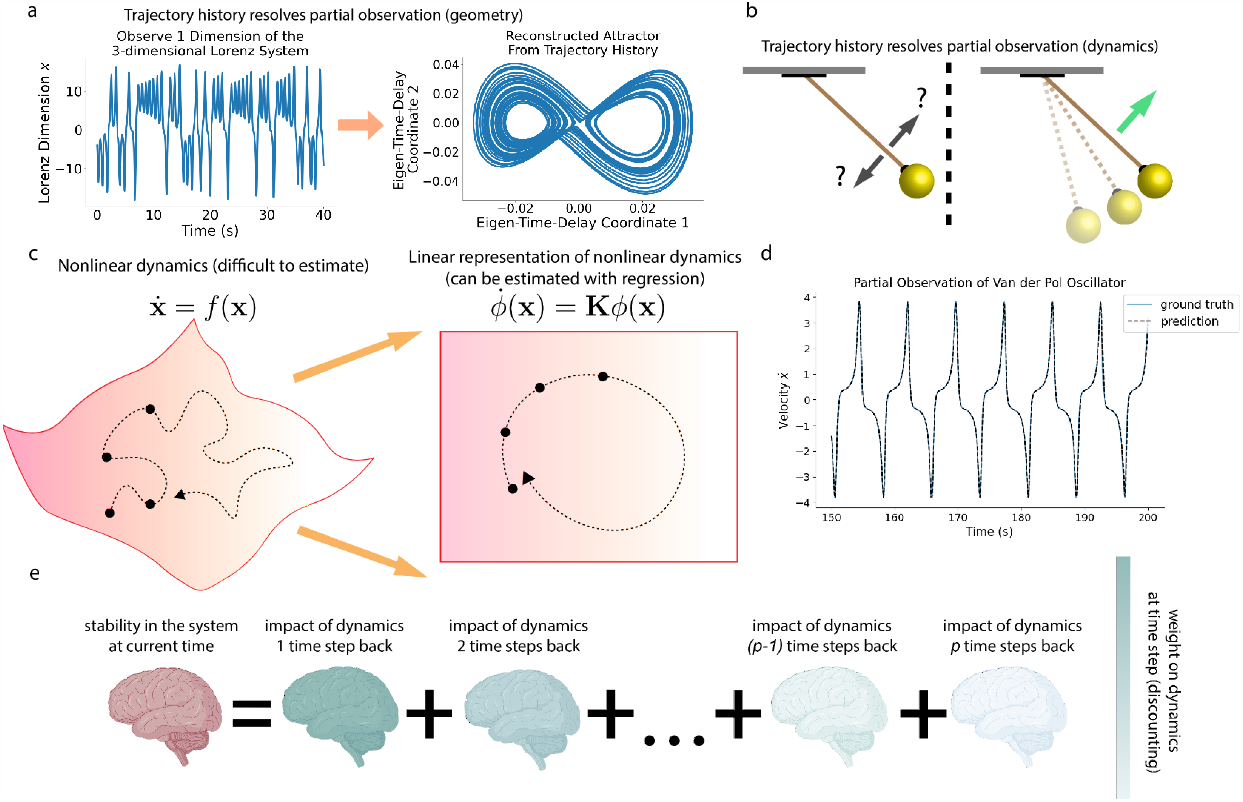
DeLASE: measuring stability through linear delayed dynamical models. (a) Trajectory history enables attractor reconstruction from partial observation. A single observation of the three-dimensional Lorenz system is shown in the left panel. With only this one observation, it is not clear what the attractor geometry is. However, by performing a delay embedding the attractor geometry can be reconstructed. In the right panel, we plot the first two eigen-time-delay coordinates of the data, equivalent to performing principal component analysis (PCA) whitening on the delay embedding matrix. This reveals that the delay embedding has reconstructed the famous butterfly attractor. (b) Trajectory history enables prediction in partially observed systems. In the left panel, a single snapshot of a pendulum is shown - with only the position, no prediction can be made. In the right panel, the trajectory history illuminates the impact of the unobserved variable (the pendulum velocity) on the observed variable (the position). (c) A cartoon depiction of the Koopman operator. A flow on a nonlinear manifold can be expanded (in this case through time-delay embedding) into a high-dimensional embedding in which there exists a linear representation for the dynamics. (d) Dynamical modeling of the partially-observed second dimension of the nonlinear Van der Pol oscillator. The HAVOK models are able to achieve a fully autonomous (i.e. future predictions are generated by previous ones) linear representation of the nonlinear oscillator. (e) A cartoon of the approach for assessing stability from delay differential equations is illustrated. The impacts of modeled dynamics are weighted based on how far back they are from the current state.

The trajectory history of a few variables can not only construct the full attractor of a system, but also reconstruct the time-resolved dynamics from partial observations^47–53^. For instance, if we observe a snapshot of the pendulum’s position, we cannot predict its next state (Fig. 2b). As soon as we include partial observations from the trajectory history, the upcoming dynamics of the pendulum become clear. We harness the information provided by the trajectory history of neural data in a similar way to predict future states.

### Tractability and nonlinearity

There are many efficient tools for accurate prediction of linear systems. However, neural and neural circuit dynamics are highly nonlinear, and nonlinear prediction is often both challenging and computationally expensive. Here we leverage a second deep insight from dynamical systems, the Koopman operator theory, which shows that a *nonlinear* dynamical system can be represented without approximation as an *infinite dimensional linear* system^54–56^ (Fig. 2c). The major challenge in practically exploiting this theoretical insight is to find large but *finite*-dimensional representations that allow the Koopman operator theory to approximately hold.

Given that incorporating trajectory history through delay embedding reconstructs the underlying attractor from partial observation, we might surmise that the dimension-expansion from delay embeddings constitutes a reasonable finite-dimensional representation in which the Koopman theory approximately holds^35,36,40,41,57–59^. An approach that exploits exactly this insight is the Hankel Alternative View of Koopman (HAVOK)^35^, which uses a de-correlated and low-rank representation of the delay embedding matrix (known as eigen-time-delay coordinates) as an embedding space in which to estimate the Koopman operator. Under certain conditions, this approach has been shown to find an optimal finite-dimensional space for representing the Koopman operator^36^. HAVOK is a variant of Dynamic Mode Decomposition (DMD)^60–62^, an approach to estimating the Koopman operator which has been explored in many varieties^57,60,63–72^, including applications to neuroscience^41,72–76^. To demonstrate the predictive power of HAVOK models, we take a partial observation of the Van der Pol Oscillator (a two-dimensional nonlinear system)^77^. HAVOK models are able to autonomously reproduce this nonlinear time series with purely linear dynamics (Fig. 2d).

Therefore, we use HAVOK to construct efficient and accurate dynamical models of partially observed neural circuits, resolving the challenges of tractability and nonlinearity.

### Estimating stability from delayed dynamical systems models

We use the accurate dynamical models constructed with HAVOK to estimate the stability of the observed dynamics. We rearrange the dynamics equation, shifting from a Koopman representation in eigen-time-delay coordinates to a delay differential equation in the original neural space. Delay differential equations make explicit the dependence of the future states of the systems on past states. The stability of a system described by a delay differential equation is determined by the roots of its corresponding characteristic equation^78^.

These roots, known as characteristic roots, are complex-valued numbers. The real part corresponds to the (inverse) characteristic timescale at which the system will respond to a perturbation along a particular direction. The imaginary part is the frequency of the perturbation response. A negative characteristic timescale corresponds to a decay in response to a perturbation (stability). A positive characteristic timescale to an explosion in response to a perturbation (instability). Thus greater inverse timescales correspond to more instability. Note especially that given a negative inverse timescale, greater values (i.e. smaller magnitudes) correspond to *longer* timescales, because of the inverse. Thus for an overall stable system, more instability corresponds to *slower* responses to perturbations.

To numerically approximate a finite portion of the (infinitely-many) roots of a given delay differential equation, we harness the TRACE-DDE algorithm^79^. This algorithm broadly estimates stability by discounting the impact of the previous time steps based on how far back they are from the current time (Fig. 2e).

DeLASE, our approach to directly estimating stability in neural data, consists of three primary steps:

1. Performing a grid search across key HAVOK hyperparameters (the size of the delay embedding, the rank of the eigen-time-delay coordinates) to identify the optimal parameters.
2. Fitting HAVOK dynamical systems models to the data.
3. Using the TRACE-DDE algorithm to extract characteristic roots from the delay differential equations representation of the models as an estimate of stability.

### DeLASE tracks changes in stability in simulated neural networks from partial observations

To validate the DeLASE procedure, we attempted to accurately predict relative changes in stability from partial observations of systems for which the ground truth stability is known.

We began by considering simple linear dynamical systems of varying stability that are driven by noise. We picked 10 different degrees of stability of such systems, all of which were relatively close to the transition to unstable dynamics (as awake and unconscious neural states are hypothesized to be). For the stability analysis, we randomly chose 10% of the dimensions of the system for partial observation and fit HAVOK models. The correlation of the mean instability estimated from DeLASE with the ground truth stability is 1 (Fig. 3a).

**Figure 3.**
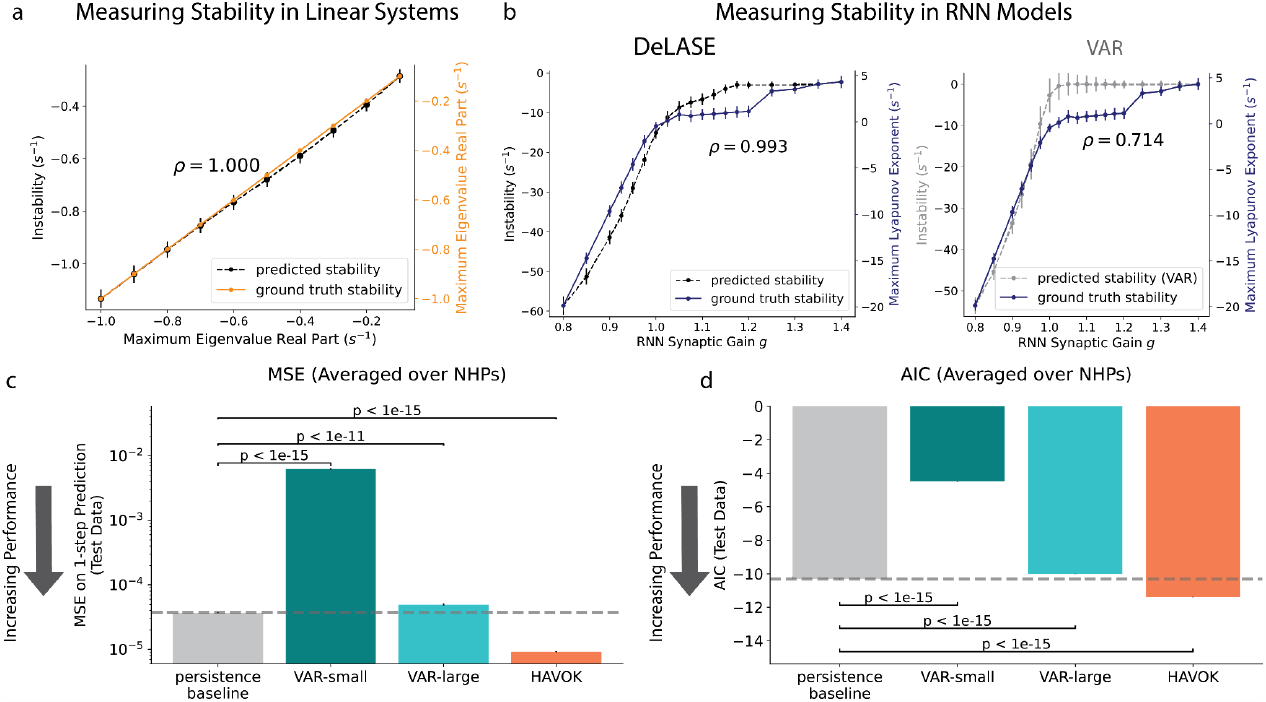
Validating DeLASE on simulated and empirical data. (a) Predicting stability in linear systems using DeLASE. The x-axis and right y-axis are the maximal (real part) of the dynamics matrix eigenvalues (orange line). The left y-axis, corresponding to the black dotted line, shows instability estimated from DeLASE, using the maximal 10% of the real parts of characteristic roots estimated from the delay differential equations analysis. DeLASE was fit to 10 out of 100 dimensions of the system and achieved a correlation of 1 in matching relative changes in stability. (b) (left) Predicting stability in RNN models using DeLASE. The x-axis is the gain parameter, which scales the synaptic weights and increases instability in the networks. The right y-axis is the maximum Lyapunov exponent of the networks (averaged over 10 simulations) which determines the stability (blue curve). The full system state and dynamics are needed to measure this in the standard approach we used^82^. The left y-axis, corresponding to the black dotted line, shows instability estimated from DeLASE, with the same approach as for (a). DeLASE was fit to 10 out of 1024 dimensions of the simulated system and achieved a correlation of 0.993 with the ground truth stability. (right) Same as for the DeLASE figure to the left, but using the VAR stability estimation approach - fit to a large temporal window. VAR was also fit to 10 out of 1024 dimensions, and the maximal 10% of the real parts of eigenvalues were used to estimate stability. VAR achieved a correlation of 0.714 in matching relative changes in stability. (c) Comparison of different model’s one-step predictions on neural data (one session from each NHP). Predictions were generated based on local field potentials recorded from multi-electrode arrays placed in four areas (approximately 250 electrodes total). Data was sampled at 1 kHz. Error bars are shown. The persistence baseline - predicting the next state as identical to the previous state - is plotted in the gray bar and also with the dotted line. HAVOK models are the only model shown capable of beating the persistence baseline at this essential prediction task. (d) The same as (c), but with AIC, depicting that the superior performance of HAVOK is not due to having more parameters than the other models.

To further validate the DeLASE procedure on more brain-like systems, we considered numerous randomly connected recurrent neural networks (RNNs) with a gain parameter . The gain parameter scales the synaptic weights and induces a transition to chaos in these networks^80^. We ran DeLASE on a randomly chosen partial observation of approximately 1% of the dimensions from each network. DeLASE predicted relative changes in the stability of RNNs with a correlation of 0.993 (Fig. 3b). For reference, we compare DeLASE to Vector Autoregression (VAR), a simple linear dynamical systems model, which has been used previously to estimate stability in neural data^31,32,81^.

## Delayed linear dynamical systems models capture neural dynamics

Good estimates of stability depend on good models of dynamics. We chose HAVOK models (see Methods) because they use a history of neural states to construct simpler linear dynamics from a nonlinear system (like the brain). Our first step towards estimating stability in real neural data, therefore, was to confirm that HAVOK was a good model for capturing neural dynamics.

We compared HAVOK to three other models. (1) A persistence baseline model. This model predicts the neural state at each time step will be identical to the previous time step. Better models of neural dynamics should thus be able to outperform the persistence baseline model. (2) + (3) Two forms of Vector Autoregression (VAR) models, which were previously used to study stability in propofol-induced unconsciousness^31,32^. Here we use VAR(1) models, which generate predictions for the next state using only the most recent state - but also take into account the dynamics across the training set. One VAR model (VAR-small) used 500 milliseconds of neural data for training, as in previous work^31^. The other VAR model (VAR-large) used a 15 second window. VAR-large was included because we used 15 second windows for HAVOK. All models were tested on a window of data of equal size to the training window, and temporally immediately following the training window. HAVOK, in contrast to VAR, predicts future states by taking into account the history of multiple preceding timesteps.

Only the HAVOK model outperformed the persistence baseline model. We compute mean-squared error (MSE) of one-step model predictions averaged over two sessions (one from each NHP) (Fig. 3c). The VAR-small models were significantly outperformed by the persistence baseline models (p < 1e-15, two-sample t-test, one sided). The VAR-large models achieved an MSE much closer to the persistence baseline but were still outperformed (p < 1e-11, two-sample t-test, one sided). Only the HAVOK models are capable of beating the persistence baseline (p < 1e-15, two-sample t-test, one sided). Akaike Information Criterion (AIC) was also used to assess the models’ prediction quality, relative to the number of parameters. AIC penalizes models for complexity and thus guards against overfitting data. HAVOK models have more parameters. Again, only the HAVOK models outperform the persistence baseline models on one-step prediction (Fig. 3d, p < 1e-15, two-sample t-test, one sided). All results held when considering the prediction quality for each NHP separately (Extended Data Fig. 1).

Thus, through the inclusion of the history of neural states, HAVOK models form accurate dynamical models of the multi-electrode activity while preserving the tractability and interpretability of linear methods.

## Propofol anesthesia induces unstable cortical neural dynamics

To determine the impact of propofol anesthesia on the stability of neural dynamics, we analyzed multi-electrode activity recorded from two non-human primates (NHPs)^2^. Electrodes were placed in four areas: ventrolateral prefrontal cortex, frontal eye fields, posterior parietal cortex and auditory cortex. We found that propofol destabilized neural activity.

We first characterized the stability of every brain region separately using DeLASE (Fig. 4a). Characteristic roots were used as a metric of instability. One component is the timescale of the system response to perturbation. The faster the response, the more stable the system. We analyzed the instability values from the upper 10% of the distribution of characteristic roots (Fig. 4a). We used the upper 10% because they have the most impact on dynamic stability. For both animals and across all four cortical regions, propofol anesthesia reliably destabilized neural dynamics (blue and gray curves, p < 1e-15 for all combinations of NHP/area with a Mann-Whitney U test). The same was true for all areas considered together as a single system (purple curves, p < 1e-15 for both NHPs with a Mann-Whitney U test). Note that for each curve, the instability values increased after propofol induction. This means the system was slower to respond (i.e., less stable).

**Figure 4.**
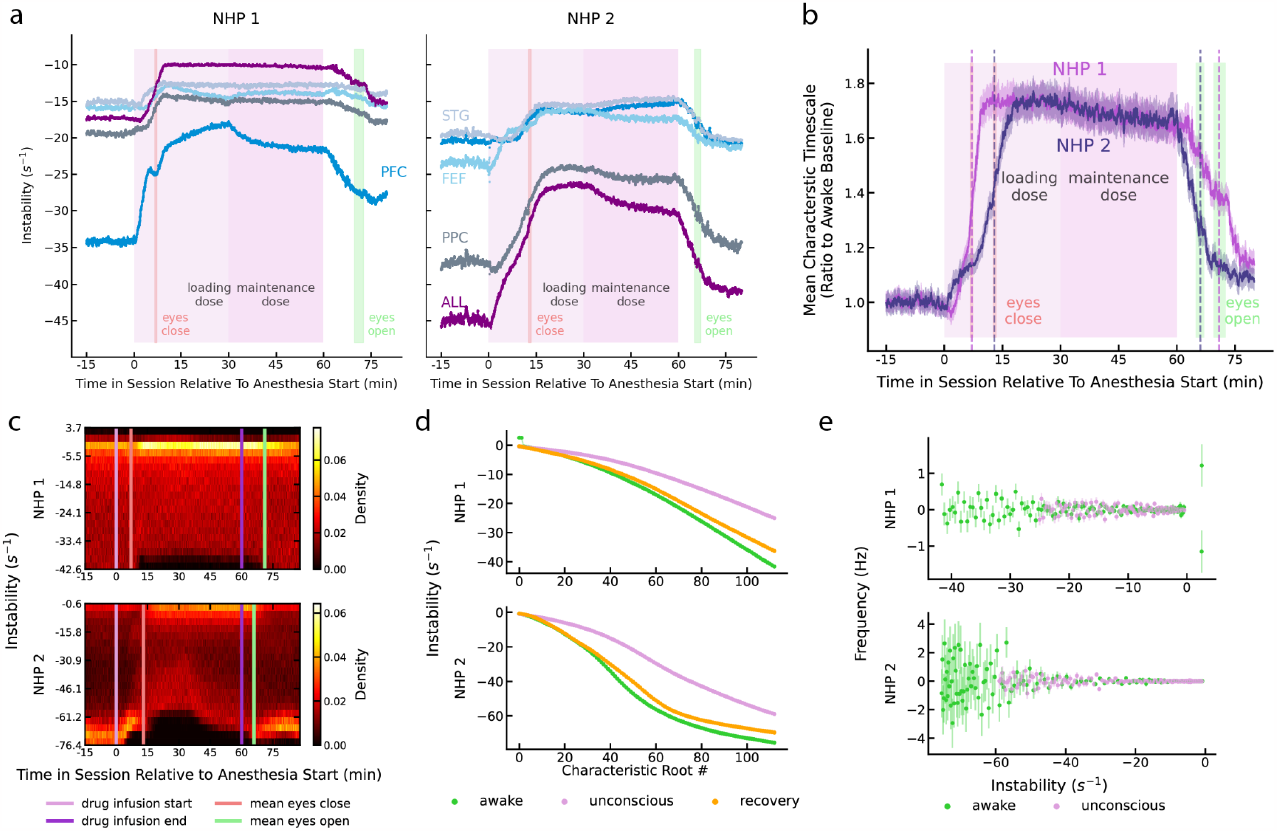
Propofol anesthesia destabilizes neural dynamics. All characteristic root analyses refer to the maximal 10% of the distribution of characteristic roots extracted from delay differential equations analysis. (a) Instability across the session in both NHPs (standard errors are plotted). DeLASE was fit to contiguous windows of 15 seconds across the session for each area individually. The x-axis is time relative to the start of anesthesia. The y-axis is the mean real part of characteristic roots (the inverse timescale of response) - a measure of instability. Instability increases during anesthesia for all areas in both NHPs. The red line indicates the mean moment of eyes closing (used as a proxy for loss of consciousness) and the green line indicates the mean moment of eyes opening (used as a proxy for return of consciousness). (b) The curves representing the mean timescales of response normalized to the awake baseline for “ALL” areas considered together from both NHPs, plotted on the same axis. The awake baseline is computed for each session by taking the geometric mean of the timescales associated with the characteristic roots across all windows in the awake section of the session. Then, for each window, the geometric mean of the ratio of the timescales to the awake baseline is computed. After accounting for the baseline awake stability, the degree of destabilization in the timescales in each NHP was very similar. Vertical dotted lines are eyes close/eyes open, with color used to indicate the mean for each NHP. (c) Visualization of the mean real parts of characteristic roots across sessions in each NHP. The x-axis is time relative to anesthesia start and the y-axis is the real part of the characteristic roots. Color represents the density of characteristic roots in a particular bin. In both NHPs, the distribution shifted upwards to be more unstable during the unconscious state. (d) The spectrum of characteristic roots. Real parts are sorted then averaged across all sessions for a particular state in the session. In both NHPs, the spectrum of characteristic roots relaxed closer to the wakeful baseline during recovery. (e) Frequency information from computed roots. The x-axis is the real part of the characteristic root and the y-axis is the imaginary part of the characteristic root (converted to a frequency in Hz).

Note that the absolute values of instability were variable across areas and NHPs. Because of the nature of the instability values, their absolute value will capture details of the differences in signals due to factors such as exact location of the electrodes relative to the neurons, electrode impedance, etc. The critical measure, therefore, is the change in values over time. When normalized to baseline values, the change in instability across areas differed in magnitude but followed similar time-courses (Extended Data Fig. 2a). When we considered all recorded areas as a single system, the ratio of change in instability values were remarkably similar across the two NHPs (Fig. 4b). For the remainder of this paper we focus on models constructed from all areas as a single system (Fig. 4b). Note that instability values during unconsciousness were nearly twice that of awake states (pre-propofol), indicating that the system’s responses to perturbation were much slower than the response during the awake state.

To better visualize the change in instability values, we plot the distribution of the top 10% of values at each time point across all areas (Fig. 4c). The entire distribution of instability values shifted upward during propofol infusion (p < 1e-15, Mann-Whitney U test). This was also true when considering the instability value distribution below the top 10% (Extended Data Fig. 2b). After eyes reopened, the distribution of values shifted down to be more stable (p < 1e-15, Kolmogorov-Smirnov test), approaching that seen pre-propofol (Fig. 4d, p < 0.005 for both NHPs, 1000-sample permutation test on the Cramér-von Mises criterion of awake-recovery distributions compared to awake-unconscious).

Next, we considered another measure of system response to perturbation, the frequency of the system response (Fig. 4e). During unconsciousness, a larger portion of the frequency components fell in the lower frequency bands than during the awake state (Extended Data Fig. 2c, p < 0.05 for both NHPs, permutation test on theta for NHP 1 and delta for NHP 2). Conversely, during the awake state, more fell into the highest populated frequency bands (Extended Data Fig. 2c, p < 0.05 for both NHPs, permutation test on beta for NHP 1 and gamma and NHP 2). This is consistent with observations that propofol causes a large increase in lower frequency power and decrease in higher frequency power in cortex^2^.

## Unstable dynamics explain sensory responses in anesthetized cortex

In the previous section, we established that propofol measurably destabilizes cortical neural dynamics. We now investigate how this destabilization changes sensory responses. We found that sensory responses under anesthesia are consistent with destabilized linear filters.

A hallmark of more stable systems is that when perturbed, they recover faster than less stable systems. An example of this is two systems in which spheres of different mass are hanging at rest from identical springs (Fig. 5a). Less mass causes the system to be more stable. Thus, when perturbed with identical force, the spring with the less mass decays back to its resting state more quickly.

**Figure 5.**
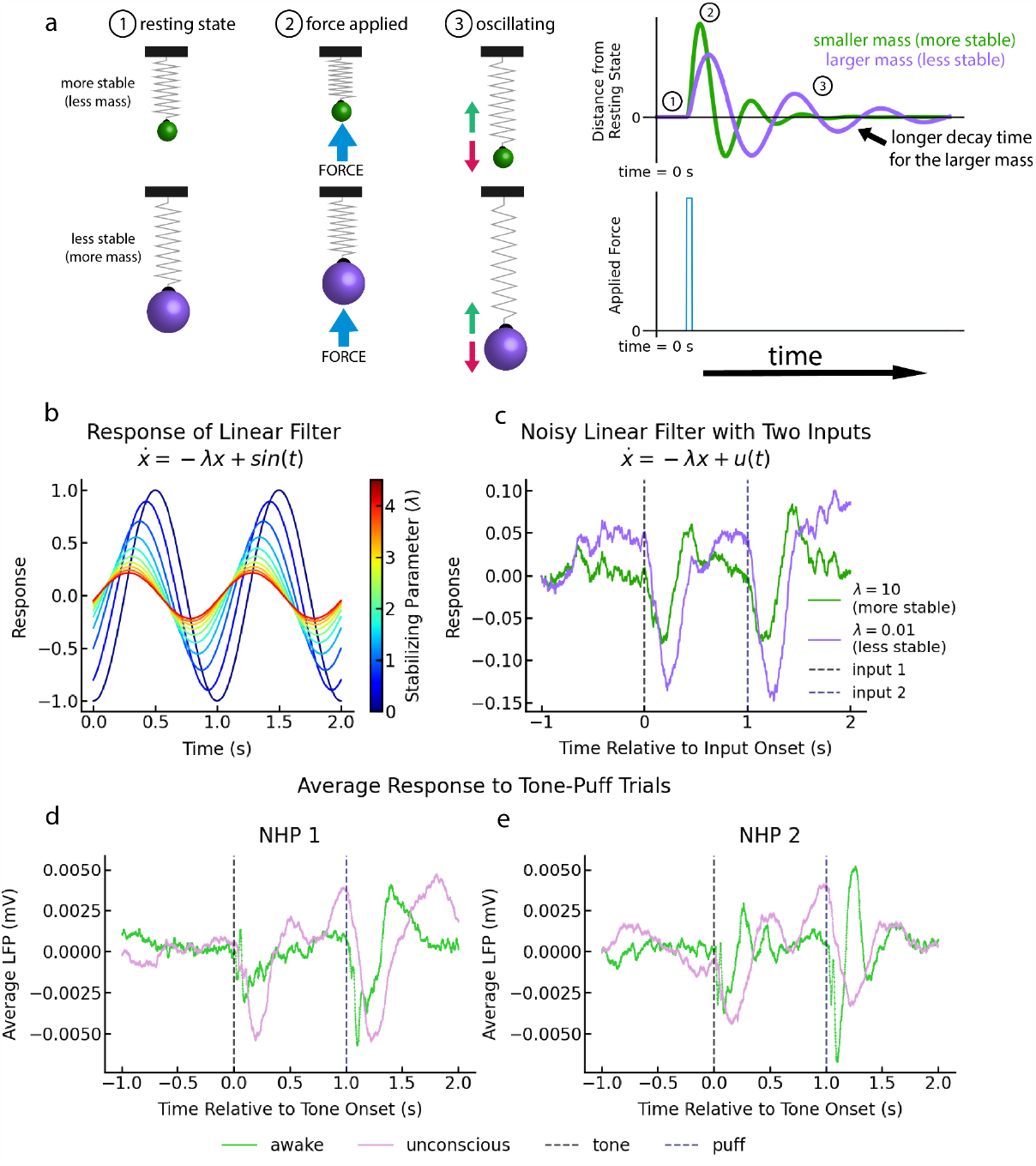
Destabilized linear filters explain responses to tones and puffs auditory cortex during anesthetic unconsciousness. (a) Damped mass-spring oscillators. An example of changes in the timescale of response to perturbation in more and less stable systems. Spheres of different mass are hanging at rest from identical springs (1). The spheres are perturbed upwards by a force of equal magnitude (2) which sets about an oscillating return to the resting state (3). The system that includes the green sphere of smaller mass is more stable and thus decays back to rest more quickly than the system that includes the purple sphere of larger mass. (b) Simple simulations demonstrating how increasing the rate of decay of the linear filter - thus stabilizing the filter - generates a phase shift and amplitude decrease in a filter driven by an oscillatory input. (c) The response to two oscillatory inputs and process noise for two particular choices of the decay parameter of the linear filter. The more unstable parameter (purple) shows an amplitude increase and phase shift relative to the less unstable parameter (green). To mimic the neural responses, two inputs were provided at times in the simulation similar to the relative timing of tones and puffs in trials. (d, e) The mean tone-evoked LFP response for all sessions from each NHP (NHP 1 is shown in (d) and NHP 2 is shown in (e), standard errors are plotted). Time relative to the tone onset is plotted on the x-axis with LFP mV plotted on the y-axis. Responses from the anesthetic unconscious state show a phase shift relative to the mean awake state response, qualitatively matching responses of destabilized linear filters.

We can describe the pattern of response to perturbation with a simple linear filter of the form 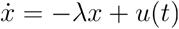, where *u(t)* is a perturbation. In this filter, the parameter λ controls the “stability” of the filter - for larger values of λ, the filter is more stable in that the filter will take less time to recover in response to perturbations. We considered the response of such a filter to a pure sinusoidal input (*u(t) =* sin*(t)* ) across a wide range of λvalues (Fig. 5b). Smaller values of λ correspond to larger amplitude, phase shifted responses. To illustrate such responses in a more neural-like setting, we simulated two different linear filters, driven by a small amount of noise (Fig. 5c). At times 0 and 1 (intended to match the timing of tones and puffs), sinusoidal inputs were provided to the linear filter. The purple curve, with a less stable parameter λ shows a response with a phase shift relative to the green curve, with a more stable parameter.

This simple model perfectly captures the qualitative change in neural responses to stimuli from the awake state to the unconscious state. Throughout the entire session, tone and puff stimuli are delivered to the NHPs (see Methods). In our analyses, we focus on trials consisting of a 500 ms tone and brief puff combination, spaced 500ms apart. We computed the mean cortical LFP response to the tones and puffs from all sessions (Fig. 5d,e). There is a phase shift in the sensory responses under anesthesia and the responses are slower than during the awake state. This exactly matches the intuition from the simulated systems: more stable systems decay back to rest more slowly after being perturbed.

## Sensory-evoked trajectories are consistent with destabilized dynamics under anesthesia

We already demonstrated that sensory responses to perturbations are slower in unconsciousness compared to the awake state. We now consider how this destabilization manifests in the neural state-space. To gain some intuition for what we might expect, we consider what happens when overall stable systems with different timescales of response are perturbed (Fig. 6a). In a system with more stability, a perturbation may cause a brief divergence from a baseline region of state-space (left). In a system with less stability, however, the perturbation induces a prolonged rotational divergence from the baseline region, depicted with slowly decaying oscillations (right).

**Figure 6.**
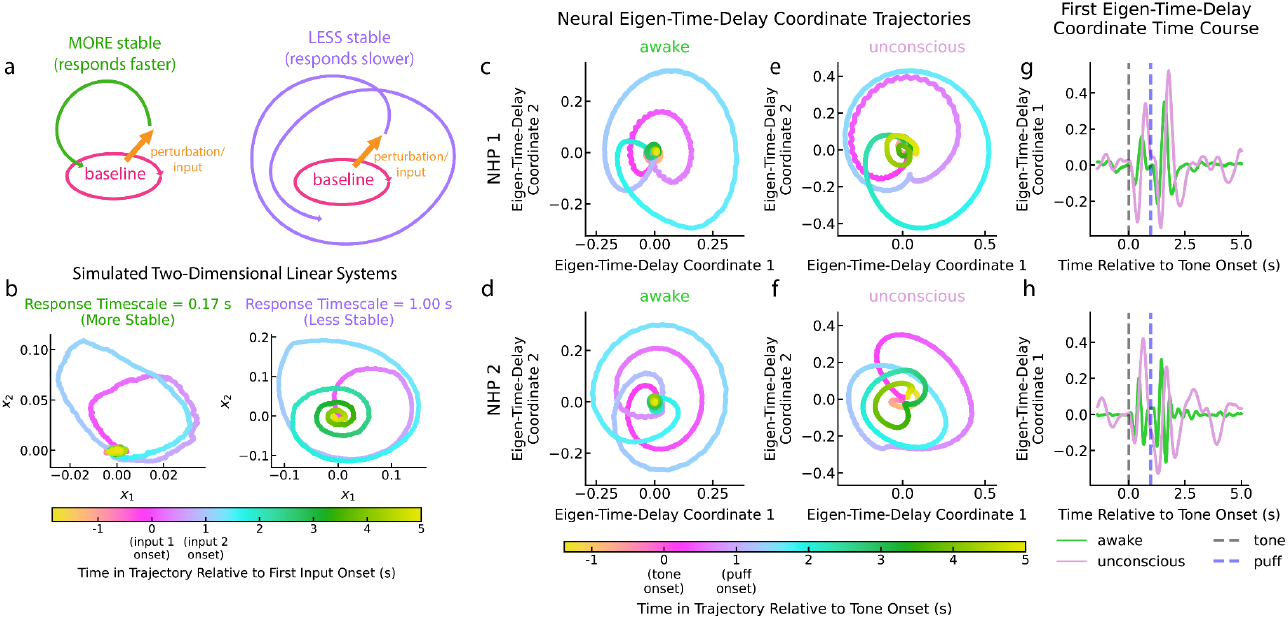
Low-dimensional trajectories of input responses in awake and unconscious states. (a) A cartoon depiction of dynamic responses to an input or perturbation when the overall system is stable. Without perturbation, the system stays within a small region of state-space (pink oval). The system is then perturbed from its baseline regime (orange arrow). In a more stable system, the system will recover quickly back to the baseline regime (left, green arrow). In a less stable system, the system will take longer to recover, potentially exhibiting decaying oscillatory dynamics on its path back (right, purple arrow). (b) A two-dimensional noise driven linear system is simulated with two different intrinsic timescales of response. In both systems, the system is perturbed with a 500 ms input at time 0 and an 150 ms input at time 1, approximately matching the structure of the inputs provided to the NHPs. The system exhibits very different responses depending on whether there is more stability (left) or less stability (right). (c, d) State-space embeddings of the mean responses to tone-puff trials from NHP 1 (c) and NHP 2 (d) during the awake state. Neural responses from tone-puff trials were each delay embedded to illuminate the attractor structure and then averaged. PCA was performed on the mean delay embedded trajectories for visualization (these are equivalent to eigen-time-delay coordinates, but scaled by the singular values - i.e. non-whitened). The tone is presented for 500 ms at time 0 and the puff is presented for about 10 ms at time 1. Awake trajectories depict quickly decaying responses to perturbations to the system dynamics due to the stimuli. (e, f) State-space embeddings of the mean responses to tone-puff trials from NHP 1 (e) and NHP 2 (f) during the unconscious state. The embedding procedure was the same as for the awake state. In contrast to the awake state, the perturbations to the system dynamics due to the stimuli decay back to the baseline region with a slower oscillatory structure. (g, h) The first PC from the awake and unconscious state tone-puff responses for NHP 1 (g) and NHP 2 (h). The unconscious response is slower in comparison to the awake response.

We simulated two two-dimensional linear systems with different levels of stability (Fig. 6b). At times 0 and 1, the system is perturbed with constant inputs, matching the structure of the tone and puff stimuli presented to the NHPs. As suggested in the cartoon, the more stable system (shorter governing timescale) is perturbed and then quickly returns back to the baseline. In contrast, the less stable system (longer governing timescale) responds to perturbations with slowly decaying oscillations.

We performed dimensionality reductions for subspace coding on the LFPs from all brain areas collected during both the awake and unconscious states. Our dimensionality reduction approach involved first performing a time-delay embedding on neural data from tone-puff trials from all sessions (we use a delay interval of 20 milliseconds and 32 delays). As discussed, delay embedding can help with attractor reconstruction when the available data constitutes a partial observation from a higher dimensional system. We then averaged each delay embedding coordinate across trials before performing principal component analysis (PCA) to obtain a visualization of neural trajectories in two dimensions. This is equivalent to computing (scaled, i.e. non-whitened) eigen-time-delay coordinates.

We visualized two-dimensional state-space trajectories of neural responses to tone-puff trials in eigen-time-delay coordinates (Fig. 6c-f). During the awake state, the state-space trajectories display two clear responses to the two stimuli: the response to the tone, at time 0, followed by the response to the airpuff at time 1 (Fig. 6c,d). These perturbations caused a deviation from the baseline region into other regions of state-space. The system quickly recovered from the perturbation, returning to the baseline state. By contrast, in unconsciousness, the perturbations due to tones and puffs caused prolonged deviations into state-space (Fig. 6e-f). These state-space trajectories are characterized by a slow oscillatory decay back to baseline, just as in the simulation. Furthermore, in the awake state, after being first perturbed by the tone, the system was able to decay back to the baseline region before the puff. However, in unconsciousness, the system did not recover from the tone before the puff was delivered. To highlight the different timescales between states, we also visualized the time course of the first eigen-time-delay coordinate (Fig. 6g,h). The unconscious curve depicts a slower neural response to stimulus perturbation, as suggested by the results from the previous section. The results of this section were consistent across variations in the delay embedding parameters (Extended Data Fig. 3).

## Increasing inhibition in random recurrent neural networks destabilizes them

We now propose a simple mechanism through which propofol can induce destabilization in neural circuits: increased inhibition.

Propofol is known to act as an agonist at GABA_A_ inhibitory receptors, thus increasing inhibitory tone. To model the effects of propofol on neural circuits, we alter the connectivity of the randomly connected RNNs (see Methods). These RNNs include a gain parameter *g* that scales the synaptic weights and induces a transition to chaos in these networks^80^. While it is therefore known that increasing the magnitude of *all* the network weights leads to a destabilization in these networks, the effects of changing *only* the inhibitory weights are unknown (Fig. 7a,b).

**Figure 7.**
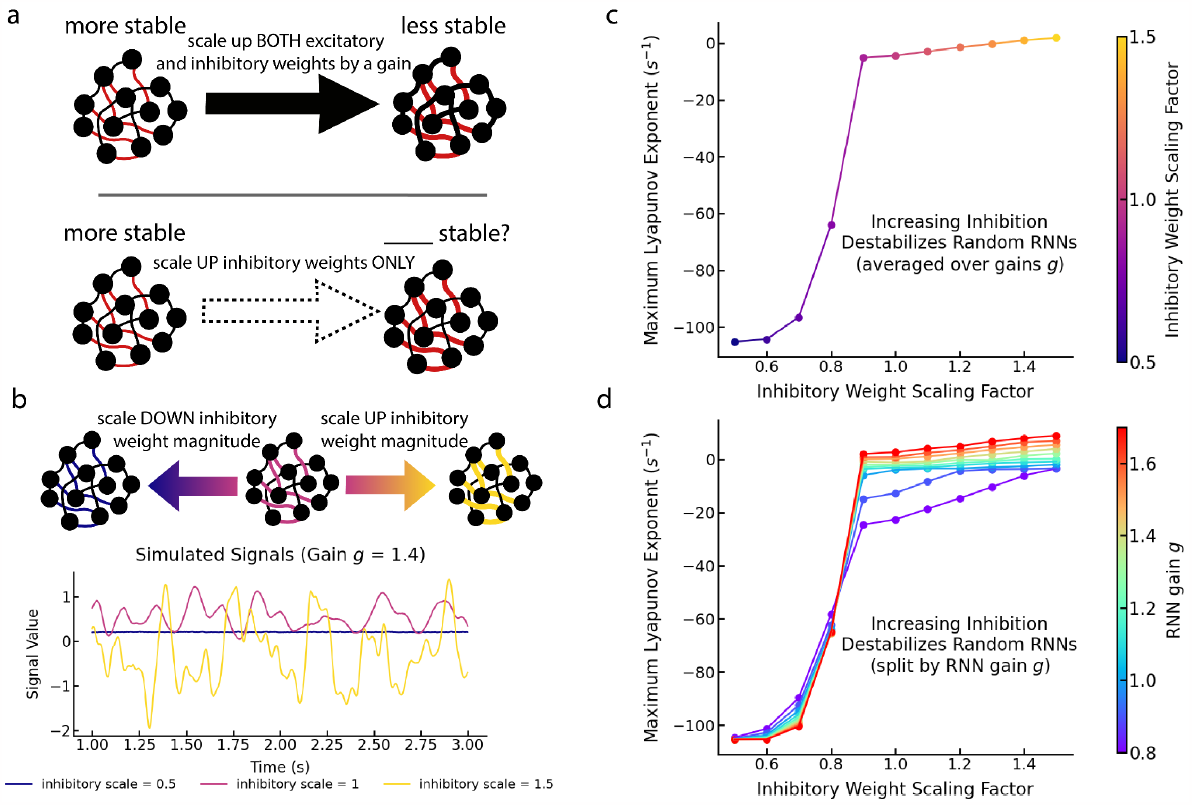
Increasing inhibition in random recurrent neural networks destabilizes them. (a) While it is well known that increasing the magnitude of all connections in random RNNs is destabilizing, it is unknown what the effect of scaling only the inhibitory weights is. (b) A demonstration of the impact of both scaling up and scaling down the magnitude of the inhibitory weights in a single RNN driven by a small amount of noise. Scaling the weights up leads to high amplitude dynamics while scaling down the inhibitory weights drastically reduces the amplitude of the dynamics. (c) The mean instability of the RNNs, as measured by the maximum Lyapunov exponent of the networks. For many different baseline gains on the connectivity weights, the RNNs were simulated 10 times. Instability was measured for each RNN. The plotted values are averaged over both simulations and baseline gains for a fixed scaling factor on the inhibitory weights. Scaling up the inhibitory weight magnitude destabilized the networks, while scaling down the inhibitory weights stabilized them. (d) The same as (c), but now split by the baseline gain. Again, we found that increasing inhibitory weight magnitude destabilizes the networks, while decreasing it stabilizes them.

We tested the impact of scaling the magnitude of the inhibitory weights in networks of varying baseline gain (thus impacting the baseline stability of the networks). We simulated the RNN dynamics with a small amount of noise (see Methods). We then measured the network instability through the maximum Lyapunov exponent, computed using standard methods^82^. We found that across all baseline connectivity gains, increasing the inhibitory tone in the networks destabilized them (Fig. 7c,d). Furthermore, decreasing the inhibitory tone in the networks stabilized them. These results were statistically significant when evaluated with a two-sample t-test (Extended Data Fig. 4).

We emphasize that this is a somewhat surprising result, given that inhibitory connectivity is often thought of as suppressing activity. In fact, it has been suggested that the increased inhibitory tone during propofol anesthesia might shut off regions of the brain during unconsciousness^6,83^.Nevertheless, the demonstrated destabilization of RNNs through increased inhibitory connectivity lends support to propofol’s action at GABA_A_ receptors underlying the observed destabilization and loss of consciousness during propofol anesthesia.

## Discussion

Our results show that propofol anesthesia destabilizes cortical neural dynamics. We found that the stability of brain dynamics is an excellent marker for anesthetic depth, as it smoothly and monotonically varies with the depth of anesthetic state. We then examined neural responses to sensory inputs. We found a longer timescale for recovery from perturbation in unconsciousness, like that seen in destabilized linear systems. We also found that increasing inhibition (as propofol does) produced destabilization. Propofol is known to disrupt the balance between cortical excitation and inhibition. Combined with our findings, this paints a picture in which propofol tampers with this balance - known to be critical for maintaining the stability of cortical dynamics^84^. The disruption of this balance causes widespread cortical instabilities, and thereby disrupts the capacity for information processing. Overall, our analysis suggests a mechanism for anesthesia that involves destabilizing brain activity to the point where the brain loses the ability to maintain conscious awareness.

We have also demonstrated the efficacy of a novel approach, DeLASE, designed to directly estimate stability in neural data. The approach brings together multiple aspects of dynamical systems theory, including delay embeddings, Koopman operators, and delay differential equations theory. The linearity of DeLASE enables it to be tractably deployed on ultra-high-volume and high-dimensional neural data. Its theoretically rigorous grounding enables it to provide robust stability estimates despite challenging features of the data such as nonlinearity and partial observation.

The extended response times to stimulus changes may contribute to the loss of conscious awareness during anesthesia. We discovered that during anesthetic unconsciousness, these timescales were nearly twice as slow as during the awake state. Therefore, when faced with an input, such as a sensory one, the neural dynamics may not be capable of the synchronization dynamics across areas required to produce conscious awareness. This is consistent with the observation that sensory cortex responses to sensory stimuli are less affected by anesthesia than those in the higher cortex^85^. Though the signal may be present in sensory areas, the full brain - including the higher-level areas necessary for conscious perception - do not converge to a combined stimulus-guided trajectory fast enough.

We wish to emphasize the use of delay embeddings as part of our methodology for analyzing the nature of neural dynamics across states. As mentioned in the methods, delay embeddings are a widely used tool known to improve the quality of dynamics and attractor reconstruction when observations are of a much smaller dimension than the dimensionality of the true system^34,35,41,47,48,53,57,86^, as is always the case in neuroscience. Delay embedding the neural data was, in our case, not only beneficial for the estimation of dynamics, but also for the visualization of neural trajectories. Patterns in neural response trajectories emerged most clearly when the trajectory history was used to reconstruct the attractor (Extended Data Fig. 3).

Our findings support the hypothesis that neural computations are instantiated by reliable dynamics in neural state-space. Given that computations such as working memory, motor control, and decision making are instantiated by dynamical features such as fixed points and attractors, destabilizing neural dynamics would likely undermine the ability of neural circuits to perform these computations^22,24,27,28,87–93^.

While we focus on the comparison between conscious and unconscious states in this paper, we emphasize that our approach to stability estimation was completely agnostic to the nature of the state that generated the neural data. It could thus be applied to a wide variety of data from a range of states. In particular, a compelling potential application is to data from psychiatric and mood disorders. Conditions such as depression, anxiety, substance use disorder, and schizophrenia can all be characterized as having distorted thinking patterns relative to neurotypical states; distortions which have been hypothesized to arise from changes to the stability landscape^19,94–98^. Tracking stability in neural dynamics over time for individuals with these conditions could massively impact the course of treatments, as well as shed light on the mechanisms of psychedelics, a novel intervention thought to disrupt overly stable dynamics^99,100^.

## Methods

### DeLASE: Delayed Linear Analysis for Stability Estimation

#### Eigen-Time-Delay Coordinates + Dynamics (HAVOK)

Here we outline the DeLASE approach to stability estimation. We consider observed data consisting of *T* observations of *N* dimensions (i.e. a matrix in ℝ *T× N*). Given a sampling interval of Δ *t*, this corresponds to a window length of *T*Δ *t*. Following HAVOK, we construct a delay embedding, i.e. a matrix of the form

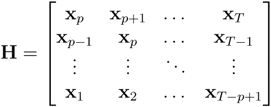

Where **x**_*t*_ ∈ ℝ^*N*^ are the state observations at time *t* (e.g. channel activities), and is the number of lags in the delay embedding matrix. Thus 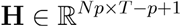 . We can now perform SVD on **H** to obtain **H** = **U ∑V**^**T**^, where **U** are the delay embedding’s spatial modes, **∑** are the singular values, and **V** are the eigen-time-delay coordinates (temporal modes). We select the *r* coordinates with largest singular values to obtain a reduced rank matrix **V**_*r*_ ∈ ℝ^*T*×^ of temporal modes. The benefits of this SVD step are twofold. First, it avoids a somewhat ill-posed regression, as a majority of the terms in each column of the delay embedding matrix *H* are identical to the previous column (but shifted). Second, it also removes any extraneous information present in the delay embedding due to correlation dimensions and noise. The computation of eigen-time-delay coordinates is equivalent to perform PCA whitening on the delay embedding matrix^75^. We then compute the dynamics matrix **A**_**V**_ ∈ ℝ^*r*× *r*^ as the matrix that solves the following least squares regression problem:

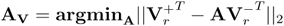

Where 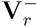 is the (*T* − *p*) × *r* matrix consisting of the first *r* eigen-time-delay coordinates at times 1 through *T* − *p* and 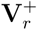 is the (*T* − *p*) × *r* matrix consisting of the first *r* eigen-time-delay coordinates at times 2 through *T* − *p+*1 . We can therefore construct a discrete model of the delay-embedded neural dynamical system as

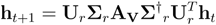

Where **h**_*t*_ is the column of the delay embedding matrix corresponding to maximal time (that is including times *t* − *p+*1through *t*), 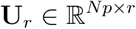 is the matrix corresponding to the first *r* columns of **U, ∑**_*r*_ is the matrix corresponding to the first *r* singular values, and **∑** ^†^ is the pseudoinverse of **∑**.

#### Estimating Stability (Delay Differential Equations Analysis)

We now let 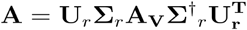 so **A** is the linear representation of the dynamics in the delay-embedded neural space. We now note that the first *N* rows of the matrix **A** describe the dependence of *x*_*t*_ on the trajectory history, explicitly

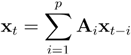

Where **A**_*i*_ is the *N* × *N* matrix corresponding to columns *iN* through (*i* + 1*)N* of the first *N* rows of **A**. We have thus arrived at a representation of the neural dynamics in the form of a (discrete) linear delay differential equation. To extract the stability from this representation of the dynamics, we turned to the field of delay differential equations. We first convert to continuous time (for purposes of interpretation) setting 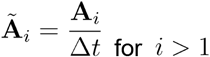 for *i* >1 and 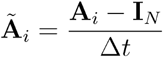 for *i* =1 (this latter term is an approximation of the instantaneous term, and so it includes an identity matrix). This yields the delay differential equation 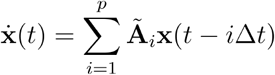 . The stability of this delay differential equation are determined by the roots λ of its corresponding characteristic equation^78^:

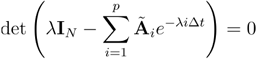

The intuition for why this equation is helpful in capturing the stability present in the system is because of the *e*^− *i*Δ *t*^ term. This term discounts the impact of the matrix **Ã**_*i*_ based on how far it is from the current state (Fig. 2e).

This characteristic equation has infinitely-many solutions^101^. In fact, even for a simple delay differential equation with a single delay, the characteristic equation has infinite solutions. Intuitively, this is because in a continuous time setting there are infinite points in between the delay time and the current time - thus the system is infinite dimensional and must be specified with an infinite dimensional initial condition. Mathematically, the infinitely-many solutions are a result of a corollary to The Great Picard Theorem, namely that an entire non-polynomial function assumes every complex number infinitely many times (with one possible exception)^102^. Analytic investigation of the stability of even single delay equations can necessitate quite advanced mathematical machinery^103^. There exist numerous methodologies to numerically approximate a finite portion of the roots of a given delay differential equation^78,79,104,105^. These approaches typically discretize the delay period, and are guaranteed to converge to the true characteristic roots as the discretization of the delay period becomes finer. For our analysis, we use the TRACE-DDE algorithm, which estimates the roots of the characteristic equation by constructing a discrete approximation of the infinitesimal generator^79^. We choose the number of discretization points in the delay period to be equal to the number of lags chosen for the delay embedding, with discretization points spaced out by Δ *t*. This generates *N*(*p*+1) characteristic roots, of which for the present analyses we evaluate the top 10% (i.e. the 10% with greatest real part).

The characteristic roots are complex valued numbers. The real part of the root determines the rate at which perturbations to the system along a particular direction will decay. The complex part of the root determines the frequency at which such perturbations will decay. In a strictly linear delay differential equation, the root with largest real part determines the stability, as it determines the overall slowest rate at which perturbations will decay (if the real part is negative) or the fastest rate at which they will explode (if the real part is positive). For our analysis, since we are approximating complex nonlinear systems with linear delay differential equations, we look at the upper portion of the distribution of characteristic roots extracted to characterize stability. Further, while the convention is to report the real parts of characteristic roots as inverse timescales, in our analysis we represent these values both with inverse timescales and standard timescales where appropriate.

To handle instabilities introduced into the dynamics by sensory stimuli, unaccounted for inputs, noise, and other artifacts, we filter the extracted characteristic roots in two ways. First we enable filtering out all characteristic roots with a frequency component above a certain value. For our analysis, we filtered out characteristic roots with a frequency component greater than 500 Hz, since this is the maximum possible frequency in data sampled at 1 kHz. Next, we enable filtering out all unstable roots with a frequency component above a certain value. For our analysis, we filtered out unstable roots with a frequency component greater than 125 Hz. The intuition for this is that rapid, sharp changes due to sensory inputs and other artifacts will destabilize the dynamics with a high frequency component, due to the sharpness of the change.

#### Picking Hyperparameters

To select (1) the minimum number of delay embedding coordinates and (2) the rank of the eigen-time-delay coordinates used to estimate the dynamics, a grid search was performed for each session (sample is shown in Extended Data Fig. 5a). We use fixed window sizes for our analysis (10 seconds for simulated data and 15 seconds for neural data). These window sizes ensure a long enough time window to capture a wide range of dynamic motifs, but a short enough time window to preserve computational tractability. For the grid search on neural data, models were fit on 12 windows from each session, and parameters were chosen on a per-session and per-area basis. Windows were chosen to ensure that all sections of the session were included (i.e. awake, anesthetic induction, anesthetic unconsciousness, and recovery), with 4 windows being randomly selected from each section in each session. The metric used to assess model quality was Akaike Information Criterion (AIC). AIC quantifies the balance between high quality model prediction (as measured by one-step prediction error on test data) with the number of model parameters. Models are penalized for having larger numbers of parameters, such that the optimal model of minimal complexity can be obtained. As can be seen in, While larger models (lower right corner of the grid) tend to perform better, after a certain point the added parameters are not benefitting the prediction quality sufficiently to justify their inclusion (Extended Data Fig. 5a). The hyperparameters minimizing AIC within a given session (and area) are chosen. The time interval between delays was held fixed at 1 time step (1 ms) for this analysis. Chosen hyperparameters varied across sessions, but were typically around 750 delay coordinates (12 delays) with a rank of 750 for individual areas (usually ∼64 channels per area), and approximately 1000 delay coordinates (4 delays) with a rank of 900 for all areas considered together (usually ∼250 channels). All tested hyperparameter combinations preserve the core results of our analysis (see Extended Data Fig. 5b).

### Implementation of VAR

We implemented VAR (VAR(1)) in the following way. We again consider observed data consisting of *T* observations from *N* dimensions (i.e. a matrix **X**∈ ℝ ^*T*× *N*^ ). Given a sampling interval of Δ *t*, this corresponds to a window length of *T* Δ *t* . We then find the matrix **A**_*VAR*_ ∈ ℝ ^*N*× *N*^ that solves the following least squares regression problem:

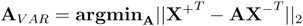

where **X**^−^ is the (*T* − 1) × *N* matrix consisting of the *N* -dimensional state observations at times 1 through *T* − 1and **X**^+^is the (*T* − 1) × *N* matrix consisting of the *N* -dimensional state observations at times 2 through .

To estimate stability, we compute the eigenvalues of **A**_*VAR*_ . To convert to a continuous time representation, for each eigenvalue λ_*I*_ we set the continuous time instability measure 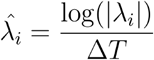 as in previous work^32^. To estimate stability, we considered the largest 10% of such instability measures extracted from VAR, mirroring our approach for DeLASE.

### Estimating Stability in Simulated Systems w/ Stochastic Dynamics

To validate the DeLASE method, and to generate examples of dynamical systems to compare to neural data, we simulated sample systems. The systems included simple linear dynamics as well as randomly connected RNNs. Since we were interested in systems that are overall stable, we often simulated these systems in the stable regime. In this regime, however, these systems converge to fixed points. Thus, to avoid this trivial behavior, we simulated these systems with stochasticity injected into their dynamics. In this approach, the systems were simulated with a small amount of process noise, effectively perturbing the system a small amount at each time step. We simulated stochastic dynamical systems using the Euler-Maruyama method. Specifically we simulated the systems using the update

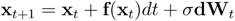

where **x**_*t*_ is the *n*-dimensional state of the system at time *t*, **f** is the system dynamics (i.e. 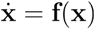, *dt* is the time step of simulation, *σ* is a small scale parameter on the noise, and **dW**_*t*_ is the *n* -dimensional process noise at time *t*, sampled from a normal distribution with mean 0 and standard deviation 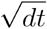 .

#### Linear Dynamics

We simulated *n* -dimensional linear systems of the base form 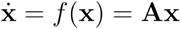 where **A** is the *n*× *n* dynamics matrix. To set the maximal real part of the eigenvalues of this matrix, we first sample each matrix element from a normal distribution with mean 0 and standard deviation 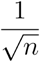 . Then we perform the update

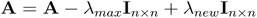

where λ_*max*_ is the original maximum real part of the matrix eigenvalues and λ_*new*_ is what we would like to update it to. We simulated the systems in the stochastic framework with *σ*=1. For the simulations used to generate Fig. 3a, we simulated stochastic linear dynamics with maximal eigenvalues from -1 to -0.1 (inclusive) stepping by 0.1. We simulated each system 20 times. Each system had 100 dimensions and was simulated for 20,000 time steps. The time step used was 0.002ms. A transient of 2000 time steps was dropped from the simulated systems, and models were fit to a randomly chosen partial observation of 10 dimensions of the system over 10,000 time steps. A hyperparameter grid search was executed to choose the number of delays and the rank of the eigen-time-delay coordinates. Specifically, minimum delay embedding matrix sizes of 10, 20, 50, 100, 200, 300, 500, 750 and 1000 were tested, along with ranks of 3, 5, 10, 25, 50, 75, 100, 125, 150, 200, all ranks from 200 to 800 stepping by 50, and also ranks of 900 and 1000. For each of the 20 runs, the best performing parameter combinations across all linear systems were chosen (typically a delay embedding matrix size of 10 with a rank of 10) and the stability was estimated based on these models.

#### Randomly-Connected RNN Dynamics

We simulated randomly-connected RNNs according to Sompolinksy et. al.^80^. Specifically, we simulated the dynamics

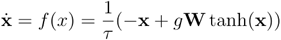

where **x** is the *n* -dimensional state of the system (effectively synaptic activity in the RNN), *τ* is the time constant of the dynamics, typically set to 100 ms (with a time step of simulation of 10 ms),**W** is the *n*× *n* weight matrix with each element drawn from a normal distribution of mean 0 standard deviation 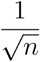, and *g* is a gain parameter on the synaptic weights. It has been shown that as *n* increases, and as the gain is increased above 1, the network enters into increasingly chaotic regimes^80^.

We used a network size of 1024 neurons, and chose values for the gain parameter from 0.8 to 1.9 (inclusive) spaced by 0.1. We sampled the matrix **W**10 times, then for each sampled matrix simulated the dynamics with all possible values of the gain. The systems were simulated for 20,000 time steps. For stability analysis, a transient of 2000 time steps was dropped, and the models were fit to 10,000 time steps of data. We randomly observed 10 dimensions of the 1024-dimensional system to use for estimating stability. A hyperparameter grid search was executed to choose the number of delays and the rank of the dynamics matrix. Specifically, minimum delay embedding matrix sizes of 10, 20, 30, 40, 50, 100, 150, 200, 250, 300, 350, and 400 were tested with ranks of 2, 3, 5, 10, 30, 40, 50, 75, 100, 125, 150, 175, 200, 225, 250, 275, 300, 325, 350, 375, and 400. For each of the 10 runs, the best performing parameter combinations across all trajectories were chosen (typically a delay embedding matrix size of 50 with a rank of 40, though there was some variation) and the stability was estimated based on these models.

To compute the Lyapunov exponents in the simulated networks, we applied a QR-based algorithm^82^ to the Jacobian of the discrete dynamics, computed as

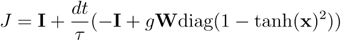

The exponents extracted from the QR algorithm are then converted into continuous time by dividing by the time step of simulation, *dt*.

### Experimental Design & Protocol

Multi-electrode neural activity was recorded from two rhesus macaques (approx. 64 channels per multi-electrode array). Electrodes were placed in four areas: ventrolateral prefrontal cortex, frontal eye fields, posterior parietal cortex and auditory cortex^2^. In the present analysis, we use LFPs from each area. There are 21 anesthetic sessions across two non-human primate subjects (NHPs). In each session, the NHP performs a non-demanding delayed saccade task pre-anesthesia. Then, following this task, the NHP experiences a passive airpuff/tone classical conditioning task. Specifically, on one third of trials the NHP is delivered a 500 ms ringing tone, which after a 500 ms blank delay, is followed by an airpuff (lasting about 10 ms) to the eye/face designed to elicit a blink response (notably, the airpuff also emits a sound). In another third of trials there is an isolated airpuff with no tone, and on the last third of trials there is a distinct tone played that indicates that no airpuff will follow. After around 15-20 minutes of airpuff/tone classical conditioning the NHP is infused with propofol anesthesia for 60 minutes. 30 minutes at a higher loading dose (0.58 mg/kg/min for NHP 1 and 0.285 mg/kg/min for NHP 2) and 30 minutes at a lower maintenance dose (0.32 mg/kg/min for NHP 1 and approx. 0.075 mg/kg/min for NHP 2). The infusion is then stopped and the NHP gradually returns to a normal awake state.

### Statistical Tests on Instability Measurements

Statistical tests referenced in the section “Propofol anesthesia induces unstable cortical dynamics” were conducted as follows. In each NHP, the real parts (i.e. inverse timescales of response) of the top 10% of characteristic roots from every window in every session were grouped according to the section (i.e. awake, anesthesia, recovery) of the session. Awake windows were collected from the 15 minutes preceding the start of the anesthetic infusion, anesthesia windows were collected from 15 minutes following propofol infusion until minutes before the end of propofol infusion (a period of 30 minutes), and recovery windows were collected from 15 minutes following the cessation of anesthesia until 25 minutes following the cessation of anesthesia.

To test whether anesthesia destabilized neural dynamics, one-sided Mann-Whitney U tests were performed on all selected inverse timescales from a given area, comparing the distributions from the awake state and anesthetic unconsciousness. To test whether the distribution of inverse timescales shifted to be more stable during recovery, we performed a one-sided Mann-Whitney U test comparing the distributions from recovery and anesthesia. To test whether the recovery distribution returned to awake levels of dynamic stability, we first constructed a test value that consisted of the awake-unconscious Cramér-von Mises criterion, subtracted from the awake-recovery Cramér-von Mises criterion. The Cramér-von Mises criterion was chosen as it constructs a non-parametric notion of distance between distributions that takes into account the whole distribution (as opposed to the widest gap, as is the case with Kolmogorov-Smirnov distance). With this construction, if the test value is negative, it suggests that the recovery distribution is closer to the awake than unconscious state, indicating a return to awake stability levels. We then conducted a 1000-sample permutation test using the constructed test value.

### Linear Dynamics with Inputs

#### One-Dimensional Linear System Responses to Sine Waves

For Fig. 5, we simulated a simple one-dimensional linear system with input according to the dynamics 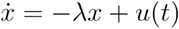 where λ controls the instability of the filter. For the simple filter on a sine wave, we found the analytical solution for the (non-stochastic) differential equation with *u(t) =* sin (2 *π ωt*). The solution is 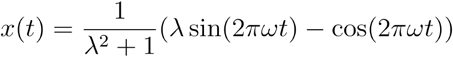 . We computed this solution for values of λ from 0 to 4.5 (inclusive), stepping by 0.5, and *ω* =1(Fig. 5b). To simulate the dynamics in the presence of noise, we simulated the simple one-dimensional stochastic dynamics

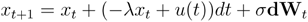

with λ set to 10 and 0.01, and *σ* set to 0.05. We used a time step of 1 ms to match the sampling rate of the neural data, and a simulation time of 3 seconds. 1 second into the simulation, we perturbed the above equation with a negative 2 Hz sine wave for 500 ms (*u(t) =*sin(4*πt* ). This matches the duration of the tone presentations in the experimental setup. We then provided no inputs for 500 ms before providing another negative 2 Hz sine wave input for 500 ms (this matches the spacing of tones and puffs in the experimental setup).

### Two-Dimensional Linear System Responses to Constant Inputs

In the two-dimensional case, we simulated the two dimensional stochastic dynamics

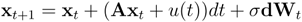

To generate the dynamics, we took two parameters, and, as well as an orthogonal matrix **Q**, and set

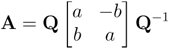

Thus **A** has eigenvalues *a* ± *bi* and *a* controls the stability of the dynamics. We simulated the dynamics with both *a* = − 6 and *a* = − 1. We set *b* = 2 *π*, and picked **Q** randomly. We used a time step of 0.25 ms and a simulation time of 7 seconds, and set *σ* =0.005. The first input was a constant function *u(t)*= 0.6 lasting 500ms and starting at 2 seconds. The second input was a constant function *u(t)*= 1 lasting 150ms and starting at 3 seconds. The structure of the inputs was chosen to mirror the tones and puffs in the experimental setup.

### Increasing Inhibition in Randomly-Connected RNNs

We again simulated randomly-connected RNNs according to Sompolinksy et. al.^80^ according to the dynamics

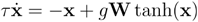

Where **x** is the *n*-dimensional state of the system (effectively synaptic activity in the RNN), *τ*is the time constant of the dynamics (10 ms, with a time step of simulation of 1 ms), is the *n* ×*n* weight matrix with each element drawn from a normal distribution of mean 0 standard Deviation 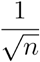, and *g* is a gain parameter on the synaptic weights. We used a network size of 512 neurons. We simulated the dynamics with gains 0.8, 0.9, 1, 1.1, 1.2, 1.3, 1.4, 1.5, 1.6, 1.7. To scale up and down the inhibition in the network, we took all elements of **W** with negative weight, and scaled them by an inhibitory scaling factor, *κ*_*I*_. For each gain, we simulated the network with *κ*_*I*_ = 0.5, 0.6, 0.7, 0.8, 0.9, 1, 1.1, 1.2, 1.3, 1.4, and 1.5. We sampled the matrix 10 times, then for each sampled matrix simulated the dynamics with all possible pairings of *g* and *κ*_*I*_. The systems were simulated for 10,000 time steps. We computed the Lyapunov exponents of the networks as described above in the section “Randomly-Connected RNN Dynamics”.

### Simulated Dynamical Systems used as Examples

#### Modeling the Van der Pol Oscillator with HAVOK

We simulated the Van der Pol Oscillator^77^ with a time step of 10 ms and a simulation time of 200 seconds (generating 20,000 time steps) using the governing equations

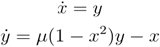

We set the parameter *μ* to 2. We then observed only the second (*y* ) dimension of the dynamics. We fit a HAVOK model on 15,000 time steps using 500 delays. Predictions were generated by autonomously running the dynamics - that is, the initial state is fed through the model to get the first prediction, then that prediction is fed back through the model, and so forth.

#### Mass-spring Simulation

For the cartoon shown in Fig. 5a, a mass-spring system was simulated with governing dynamics

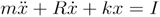

where *x* is the vertical position of the mass, *m,k,R* are parameters governing the dynamics and *I* is the input. For our simulations, we set *m* =10,*k*=1,*R*=10 . We simulated using scipy’s *solve_ivp* function for 1,000 time steps with a time step of 10ms. From time step 100 to 120, we perturb the system with a constant input of value *I* =10.

## Acknowledgements

We thank Chandrika Prakash Vyasarayani and Antonio Carlos Costa for helpful discussions and thoughts.

## Extended Data Figures

**Extended Data Figure 1.**
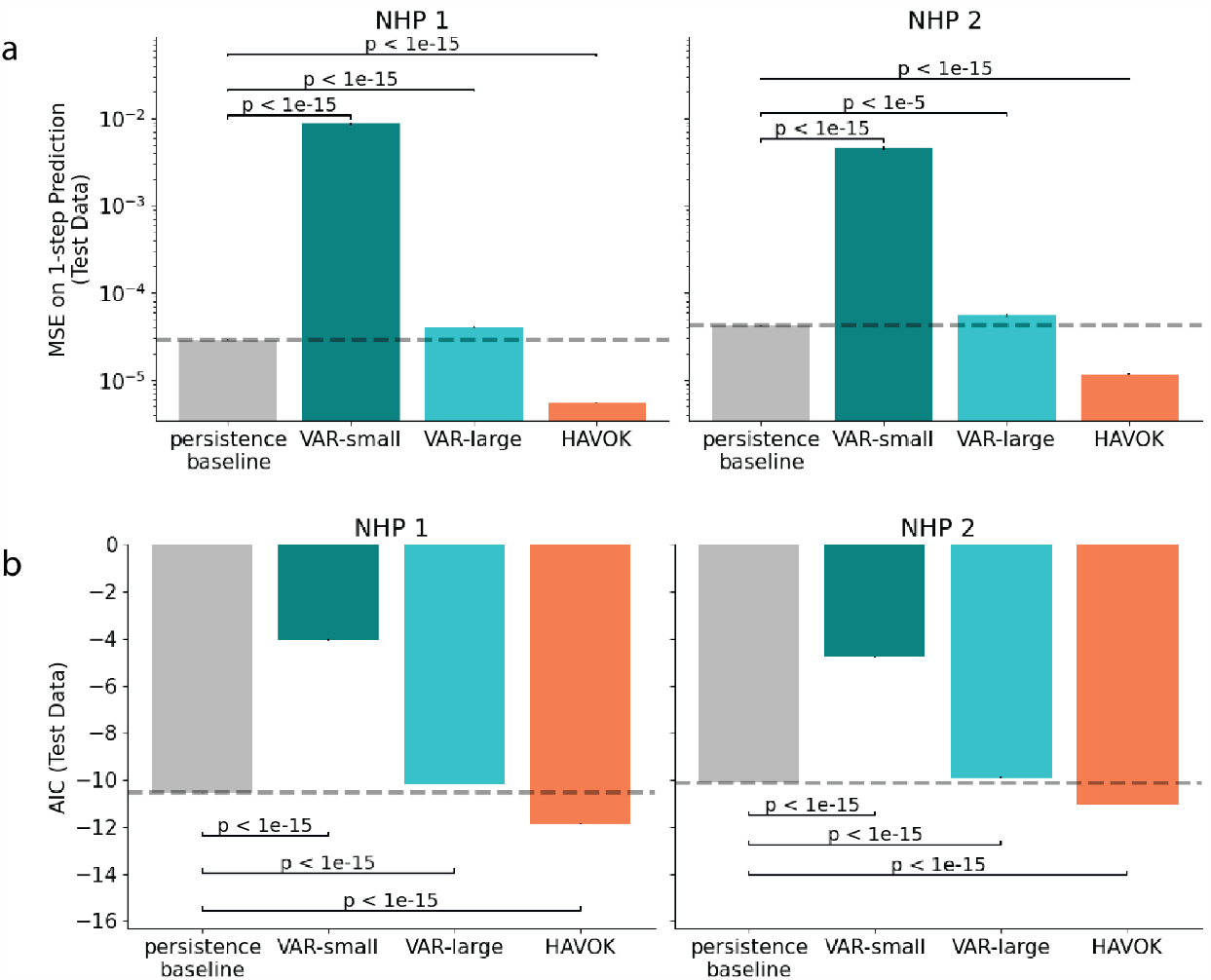
Extended methods. (a) Comparison of different model’s one-step predictions on neural data - same as Fig. 3c, but split by NHP.. The persistence baseline - predicting the next state solely with the previous state - is plotted in the gray bar and also with the dotted line. The y-axis is plotted on a log scale. HAVOK models are the only model shown capable of beating the persistence baseline at this essential prediction task. (b) The same as (a), but with AIC, depicting that the superior performance of HAVOK is not due to having more parameters than the other models. The same as Fig. 3d, but split by NHP.

**Extended Data Figure 2.**
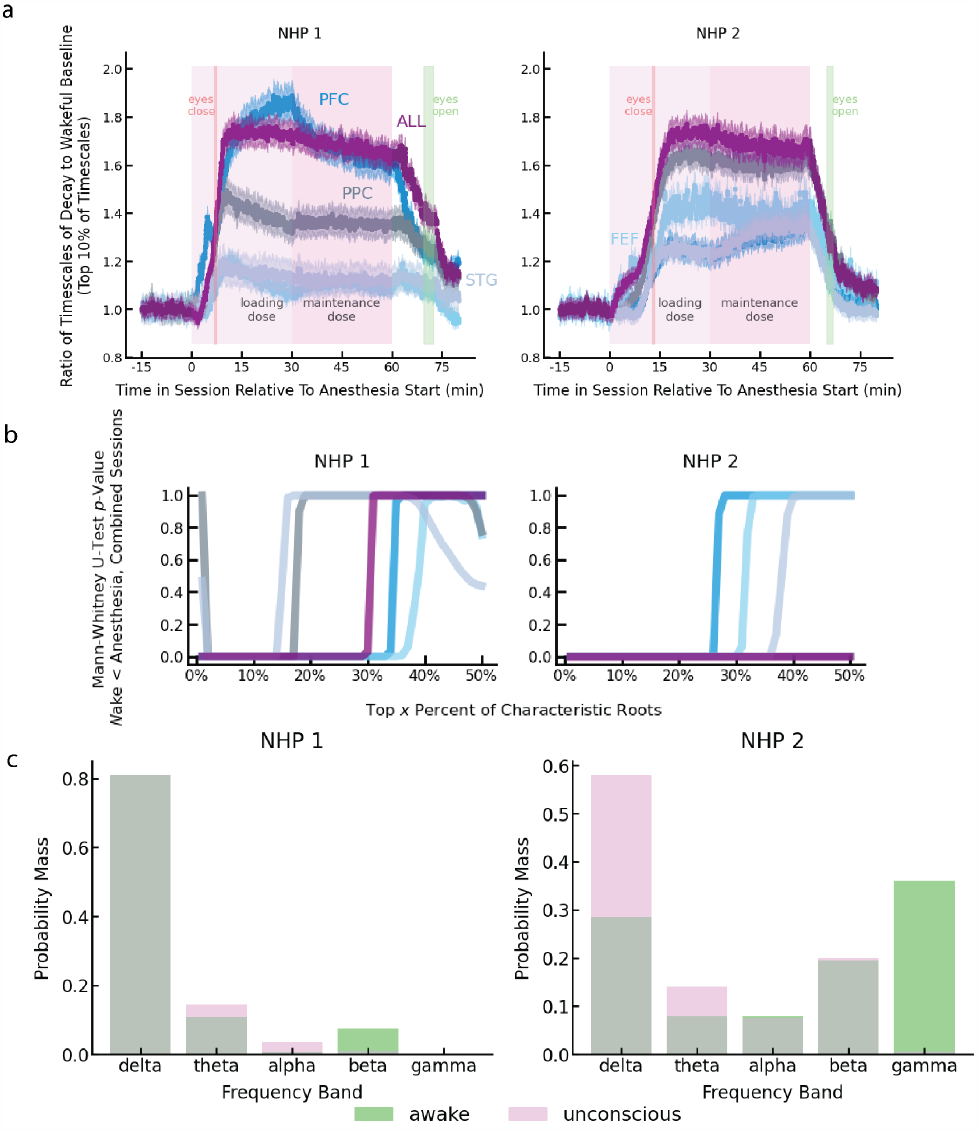
Extended neural results. (a) The curves representing mean timescales normalized to the awake baseline for all areas from both NHPs. The same as Fig. 4b, but showing results from all areas. The awake baseline is computed for each session by taking the geometric mean of the timescales associated with the characteristic roots across all windows in the awake section of the session. Then, for each window, the geometric mean of the ratio of the timescales to the awake baseline is computed. After accounting for the baseline awake stability, the degree of destabilization in the timescales in each NHP is very similar. (b) The p-value from the Mann-Whitney U-test testing whether the distribution of characteristic roots from the awake state is less than the distribution of characteristic roots from anesthesia. For each analysis, we consider the maximal x% of characteristic roots, where x is the x-axis value. For all areas, and for almost all portions of the distribution above 20%, the upper portion of the distribution of characteristic roots is more unstable in anesthesia. For higher level areas, such as vlPFC and FEF, and when considering all areas together, this is true in both NHPs for portions of the distribution greater than 30%. (c) The probability mass function on which bands the frequency components of the (maximal 10%) of characteristic roots fall into. Anesthesia (purple) is compared to the awake state (green). There is more mass on lower frequency bands in anesthesia, and more mass on higher frequency bands during the awake state.

**Extended Data Figure 3.**
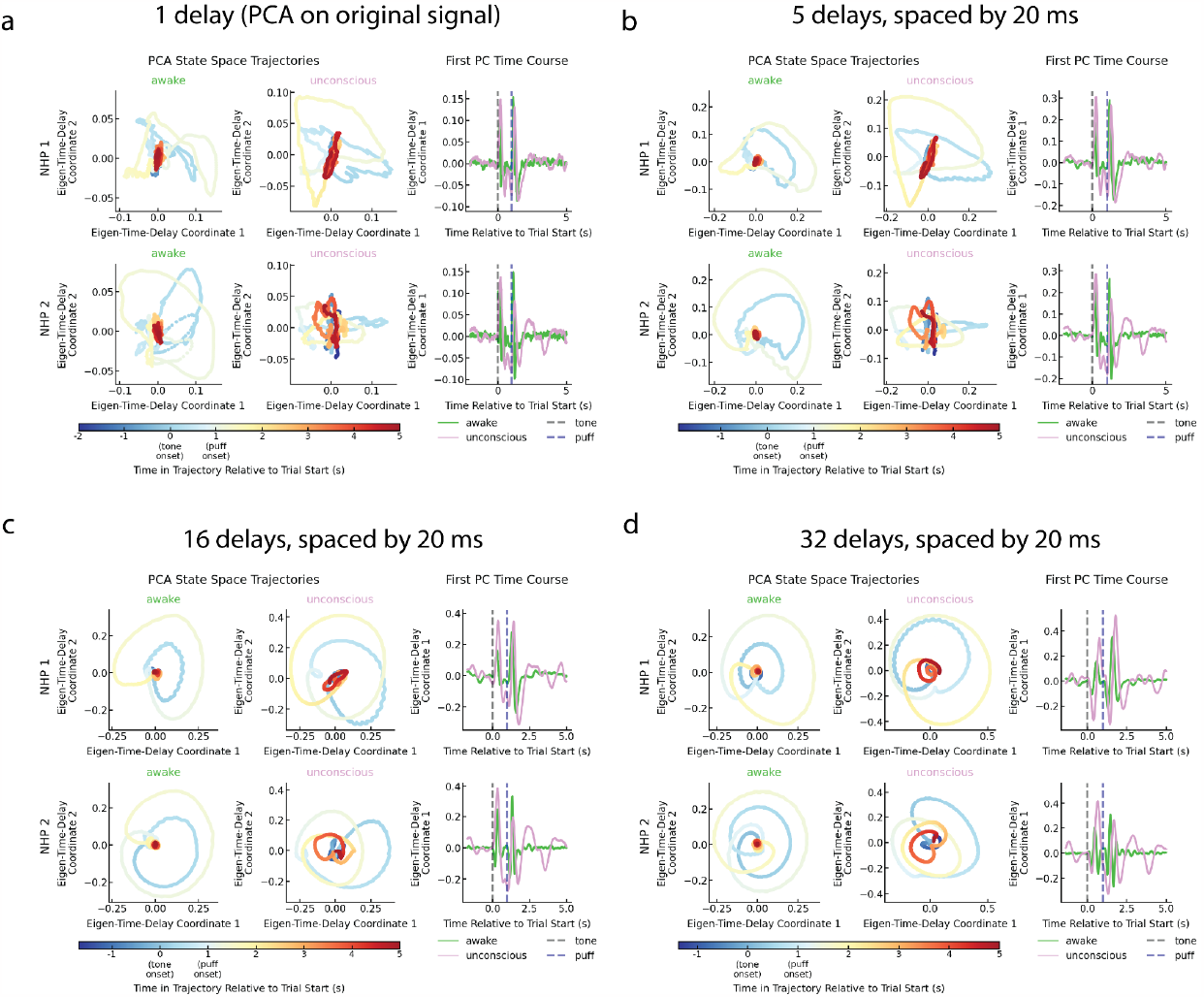
Dimensionality reduction across delay embedding hyperparameters. The core qualitative pattern is preserved across hyperparameters, though eigen-time-delay coordinates more clearly illuminate the state-space dynamics. (a) The original signal. The eigen-time-delay coordinates in this case reduce to PCA. (b) 5 delays, each spaced out by 20 ms. (c) 16 delays, each spaced out by 20 ms. (d) 32 delays, each spaced out by 20 ms.

**Extended Data Figure 4.**
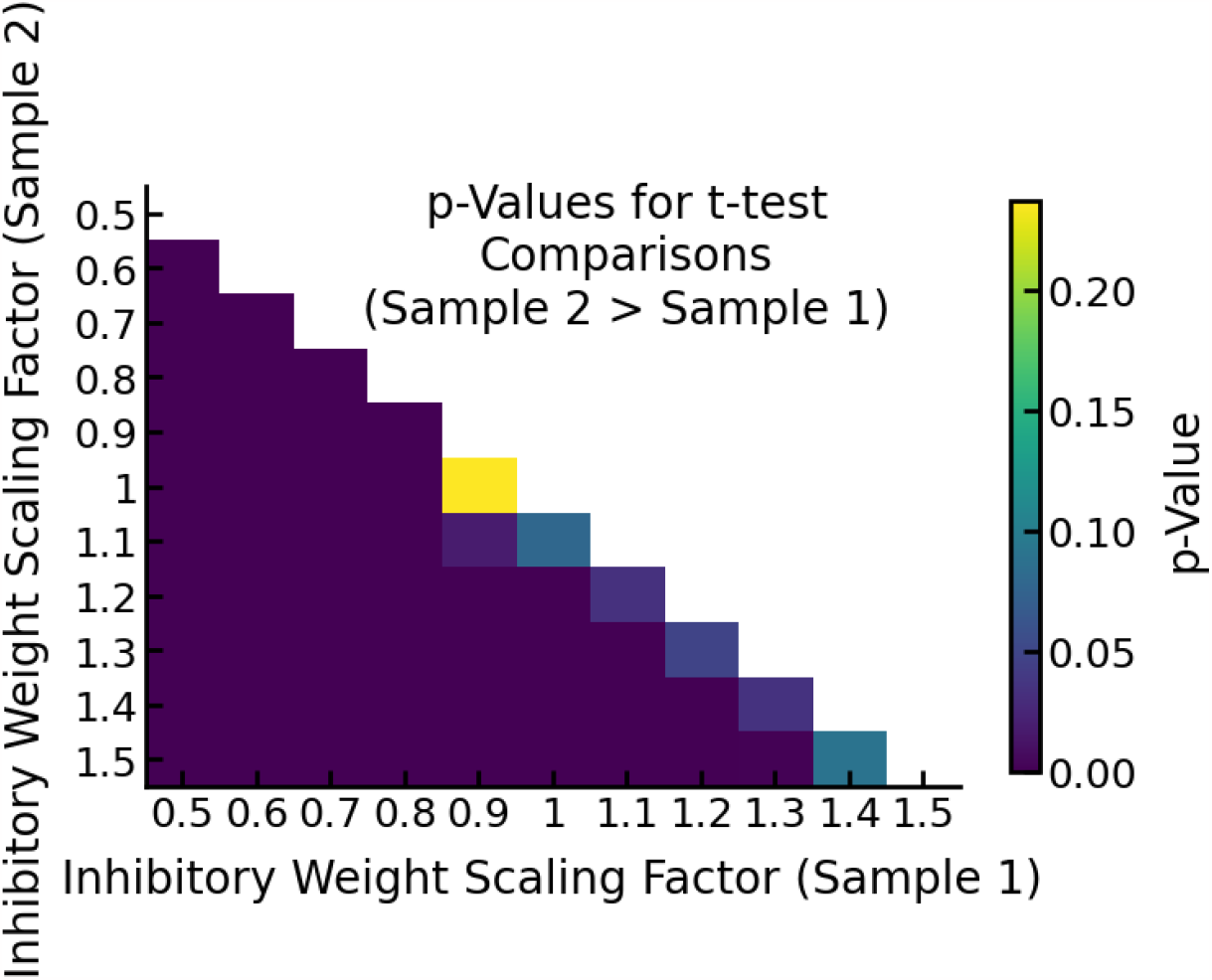
p-Values for t-test comparisons between the stability of RNNs with scaled inhibitory weights. We compared the distributions of maximum Lyapunov exponents from all the simulations at varying values of the gain parameter, which controls the stability in the networks. For each inhibitory weight scaling factor, 10 simulations were analyzed. Comparisons are across inhibitory weight scaling factors.

**Extended Data Figure 5.**
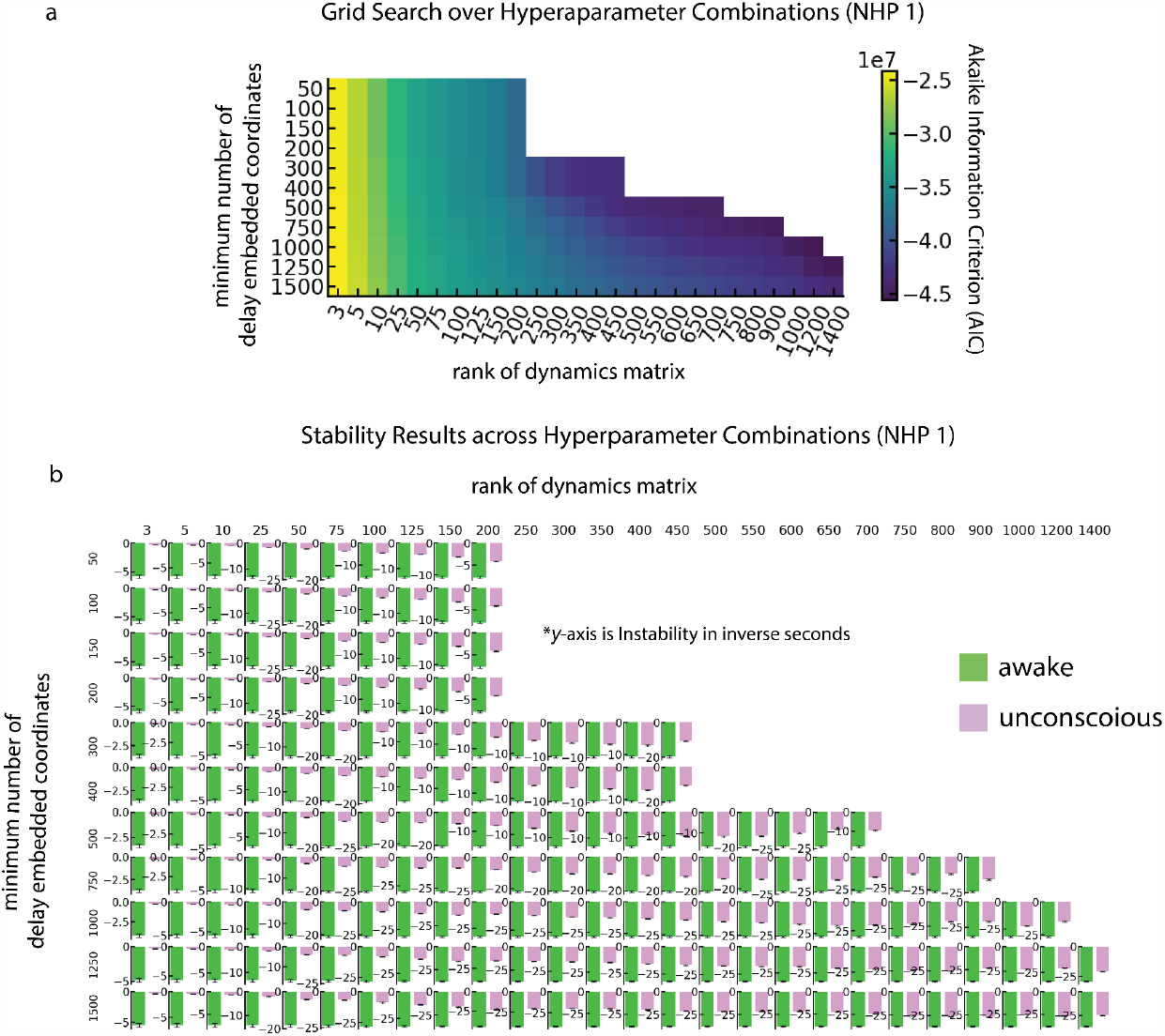
Core results hold across hyperparameter combinations. (a) The mean AIC on test-set one-step prediction on 20 randomly chosen windows of awake data and anesthetic data (10 from each) from a single session. The y-axis is the minimum number of delay embedding coordinates, effectively determining the number of lags. The x-axis is the rank of the dynamics, or number of independent temporal modes, extracted through SVD on the delay embedding matrix that are used for fitting the dynamical model. Color indicates Akaike Information Criterion (AIC) normalized to the number of predicted values. AIC quantifies the balance between high quality model prediction (as measured by one-step prediction error on test data) with the number of model parameters. Models are penalized for having larger numbers of parameters, such that the optimal model of minimal complexity can be obtained. (b) The mean of the real part of the top 10% of characteristic roots (or the number of characteristic roots equivalent to the rank, whichever is smaller) from the 10 windows from each section of the session. The core results hold across all hyperparameter combinations.

